# Soluble signal inhibitory receptor on leukocytes-1 is released from activated neutrophils by proteinase 3 cleavage

**DOI:** 10.1101/2022.03.03.482795

**Authors:** Helen J. von Richthofen, Geertje H.A. Westerlaken, Doron Gollnast, Sjanna Besteman, Eveline M. Delemarre, Karlijn Rodenburg, Petra Moerer, Daphne A.C. Stapels, Anand K. Andiappan, Olaf Rötzschke, Stefan Nierkens, Helen L. Leavis, Louis J. Bont, Suzan H.M. Rooijakkers, Linde Meyaard

## Abstract

Signal inhibitory receptor on leukocytes-1 (SIRL-1) is an immune inhibitory receptor expressed on human granulocytes and monocytes which dampens antimicrobial functions. We previously showed that sputum neutrophils from infants with severe respiratory syncytial virus (RSV) bronchiolitis have decreased SIRL-1 surface expression compared to blood neutrophils, and that SIRL-1 surface expression is rapidly lost from *in vitro* activated neutrophils. This led us to hypothesize that activated neutrophils lose SIRL-1 by ectodomain shedding. Here, we developed an ELISA and measured the concentration of soluble SIRL-1 (sSIRL-1) in RSV bronchiolitis and hospitalized COVID-19 patients, which are both characterized by neutrophilic inflammation. In line with our hypothesis, sSIRL-1 concentration was increased in sputum compared to plasma of RSV bronchiolitis patients, and in serum of hospitalized COVID-19 patients compared to control serum. In addition, we show that *in vitro* activated neutrophils release sSIRL-1 by proteolytic cleavage, which can be prevented by proteinase 3 inhibition. Finally, we found that SIRL-1 shedding is prevented by extracellular adherence protein (Eap) from *S. aureus*. Notably, we recently showed that SIRL-1 is activated by PSMα3 from *S. aureus*, suggesting that *S. aureus* may counteract SIRL-1 shedding to benefit from preserved inhibitory function of SIRL-1. In conclusion, we are the first to report that SIRL-1 is released from activated neutrophils by proteinase 3 cleavage and that endogenous sSIRL-1 protein is present *in vivo*.

## INTRODUCTION

Immune inhibitory receptors (IIRs), also referred to as immune checkpoints, are pivotal in negative regulation of immune cells (1, 2). Although far less studied than the transmembrane form, many IIRs also have a soluble form or homologue (3, 4). Soluble IIRs can arise from ectodomain shedding of the membrane-expressed receptor (5, 6), or can be a product of alternative splicing (7, 8) or a homologous gene (9). Soluble IIRs were found to be increased in the circulation of patients with several forms of cancer (reviewed by (3)), sepsis (10), and COVID-19 (11).

Signal inhibitory receptor on leukocytes-1 (SIRL-1), encoded by the *VSTM1* gene, is an IIR that is expressed on human monocytes and granulocytes in peripheral blood (12–14) and lung (15). On monocytes, but not granulocytes, SIRL-1 expression is associated with the single nucleotide polymorphism (SNP) rs612529T/C (14, 15). SIRL-1 inhibits innate effector functions such as Fc Receptor (FcR) induced production of reactive oxygen species (ROS) (14, 16, 17) and formation of neutrophil extracellular traps (NET) (17, 18). We recently showed that SIRL-1 recognizes amphipathic α-helical peptides, including cathelicidin LL-37 and *Staphylococcal* phenol-soluble modulins (PSMs) (19), as all well as several members of the S100 protein family (20), classifying SIRL-1 as an inhibitory pattern recognition receptor (21).

We previously showed that SIRL-1 surface expression on neutrophils and monocytes rapidly decreases after activation *in vitro* (16). Here, we hypothesized that activated neutrophils and monocytes shed the ectodomain of SIRL-1, thereby releasing soluble SIRL-1 (sSIRL-1). In addition, it has been described that *VSTM1* encodes the splice variant VSTM1-v2, which lacks the exon that encodes the transmembrane domain and is therefore predicted to give rise to a soluble form of SIRL-1 (22). Even though these potential sources of sSIRL-1 have been reported, presence of endogenous sSIRL-1 protein has not been demonstrated yet. In this study, we developed an ELISA to investigate the presence and release mechanism of sSIRL-1 protein.

## MATERIALS & METHODS

### Antibodies

We previously described the production of the SIRL-1 specific monoclonal antibodies (mAbs) clone 1A5 (12) and 3D3 (19). The SIRL-1 specific mAb clone 3F5 (IgG2a isotype) was produced in a similar fashion, with the exception that mice were immunized with a SIRL-1 expressing cell line instead of Fc-labeled SIRL-1 ectodomain. The 1A5 and 3F5 mAbs were conjugated with Alexa Fluor 647 (AF647) (Thermo Fisher Scientific) for flow cytometry. The 3D3 mAb was conjugated with biotin (Thermo Scientific) for the sSIRL-1 ELISA.

For antibody competition assays, peripheral blood mononuclear cells (PBMCs) were pre-incubated for 2 hours at 37°C with respective mAb clones. Subsequently, 1A5-AF647 was added to the PBMCs and incubated for 20 minutes at 4°C, followed by two washes with FACS buffer (PBS containing 1% BSA and 0.01% NaN3). SIRL-1 expression was then measured by flow cytometry.

### Biological samples

To isolate plasma for the ELISA spike experiment (Fig 1E) or for cell isolations, heparinized blood was obtained from healthy volunteers at the UMC Utrecht. For the rs612529T/C cohort, plasma and genotyping information was obtained from healthy volunteers from the Singapore Systems Immunology Cohort (SSIC) (23). For the RSV cohort, sputum, heparinized plasma and urine were obtained from infants with severe RSV bronchiolitis or control infants without infectious disease that were mechanically ventilated. These patients and the sputum isolation have been previously described (17). Briefly, sputum was collected by flushing tracheobronchial aspiration with a maximum of 2 mL normal saline. Sputum was then centrifuged 5 min at 500g to remove cells, and 30 minutes at 25000g to remove cellular debris. For the COVID-19 cohort, serum was used from SARS-CoV-2 infected patients that were hospitalized due to COVID-19 symptoms. As control, serum was obtained from healthy volunteers at the UMC Utrecht without COVID-19 symptoms. All samples were collected in accordance with the Institutional Review Board of the University Medical Center (UMC) Utrecht or the National University of Singapore.

**Fig 1.**
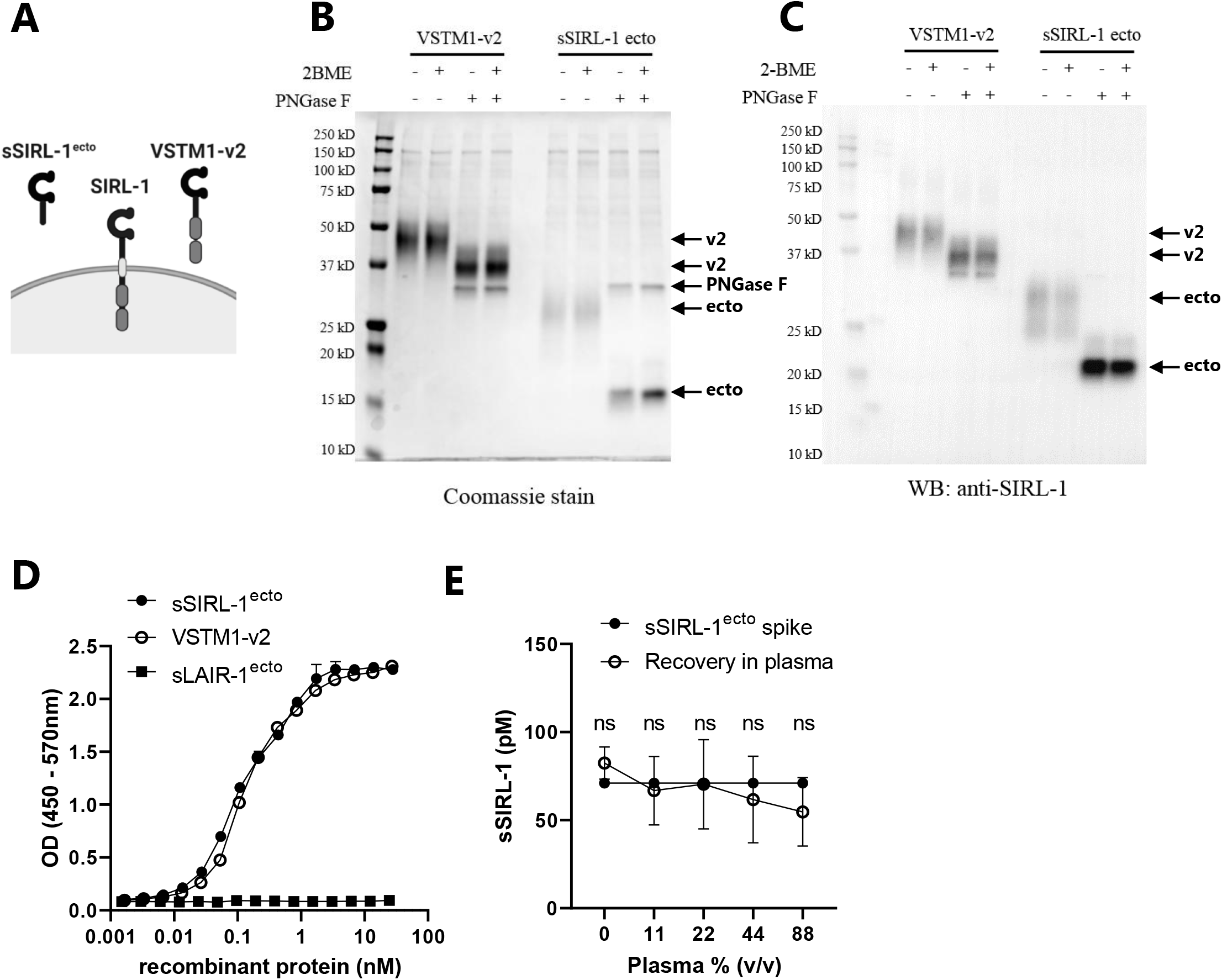
Development of sSIRL-1 ELISA. **(A)** Schematic representation of SIRL-1 and its two potential soluble forms; VSTM1-v2 and sSIRL-1^ecto^. **(B-C)** SDS-PAGE analysis of VSTM1-v2 and sSIRL-1^ecto^, with or without pre-treatment with PNGase F or β-mercaptoethanol (2-BME). Proteins were either stained directly by coomassie (B) or transferred to a membrane and stained with SIRL-1 mAb clone 1A5 (C). Arrows indicate the position of VSTM1-v2 (v2), sSIRL-1^ecto^ (ecto) or PNGase F. **(D)** sSIRL-1^ecto^, VSTM1-v2 and negative control sLAIR-1^ecto^ were titrated in an in-house developed sSIRL-1 ELISA (n≥3, one representative example is shown). **(E)** sSIRL-1 negative heparin plasmas (n=5) were titrated and spiked with 70 pM sSIRL-1^ecto^. sSIRL-1 concentration was measured by ELISA, symbols represent the mean ± SD. The statistical difference between the spike and the recovery of sSIRL-1^ecto^ at different plasma dilutions was tested using an one sample Wilcoxon test, ns = not significant.

### Protease inhibitors

The protease inhibitor cocktail (cOmplete™, EDTA-free) was purchased from Roche and used at half of the recommended concentration for lysates. Pepstatin A was purchased from Fisher Scientific, E64 from Sanbio, GM6001 from Merck, Leupeptin from Roche, Eap, EapH1 and EapH2 were recombinantly produced as previously described (24). The proteinase 3 inhibitor Bt-PYDnVP (25) (compound 10, (O-C_6_H_4_-4-Cl)_2_,) was generously provided by Dr. Brice Korkmaz and Prof. Dr. Adam Lesner and used at a concentration of 10 μM.

### Cloning and recombinant sSIRL-1 protein production

To produce cDNA constructs, gBlocks^®^ (Integrated DNA Technologies) were ordered that encode VSTM1-v2 and sSIRL-1^ecto^ with a C-terminal His-tag (see Supplementary Table 1 for the DNA sequence, and Supplementary Fig 1 for the amino acid sequence of the recombinant proteins). The gBlocks were inserted into a pcDNA 3.1+ Zeocine plasmid using Gibson assembly master mix (New England Biolabs). The cDNA construct encoding LAIR-1 ectodomain (sLAIR-1^ecto^) has been previously described (26).

Recombinant proteins were produced using the Freestyle™ 293 Expression System (Thermo Fisher Scientific) according to manufacturer’s instructions. Four days after transfection, supernatants were collected and filtered through a 0.45 μM Minisart Filter (Sartorius). The His-tagged proteins in the supernatants were purified using HIStrap FF columns (GE Healthcare) according to manufacturer’s instructions. Briefly, the columns were attached to an ÄKTAprime™ (GE Healthcare) and equilibrated with binding buffer (20 mM Na_3_PO_4_, 500 mM NaCl, 20 mM Imidazole, pH 7.4). Next, supernatants were mixed with binding buffer in a 1:1 ratio, filtered through a Stericup 0.22 μM filter (Millipore), and loaded onto the ÄKTAprime™. After loading, binding buffer was applied until a steady baseline was reached. Next, elution buffer (20 mM Na_3_PO_4_, 500 mM NaCl, 500 mM Imidazole, pH 7.4) was added for a one-step elution of the proteins. Protein-containing fractions were pooled and rebuffered to PBS using VIVAspin columns (5,000 MWCO PES; Sartorius). Protein yield was determined with a BCA protein assay (Pierce™; Thermo Scientific).

### SDS-PAGE and Western blot

To remove N-linked glycosylation, recombinant proteins were treated with PNGase F (New England Biolabs) according to manufacturer’s instructions. Next, the proteins were incubated 5 minutes at 95°C in Laemmli sample buffer (Bio-Rad laboratories), with or without addition of β-mercaptoethanol (Sigma-Aldrich Chemie). Proteins were separated by SDS-PAGE on Any kD Mini-PROTEAN TGX Precast Protein Gels (Bio-Rad Laboratories) and either stained directly with coomassie blue (Merck) or transferred onto 0.45 μm PVDF membranes (Merck) for western blot analysis.

For western blot analysis, membranes were blocked in TBS containing 0.05% Tween-20 (TBS-T) and 5% BSA, incubated with mAb 1A5 (2 μg/mL in TBS-T with 1% BSA), followed by staining with goat-anti-mouse IgG F(ab’)2-HRP (Jackson ImmunoResearch; 1:5000 diluted in TBS-T with 1% BSA). All incubations were 1h at RT and followed by extensive washing with TBS-T. Finally, proteins were visualized using ECL Western blot reagent (Fisher Scientific) and the ChemiDoc™ imaging system (Bio-Rad).

To determine PR3 cleavage, 20 μg/mL sSIRL-1^ecto^ or VSTM1-v2 were incubated with 2 μg/mL PR3 in PBS with 0.5M NaCl and incubated 3h at 25°C, followed by SDS-PAGE and Western blot analysis as described above, except that rabbit-anti-mouse IgG-HRP (DAKO; 1:10.000 diluted in TBS-T with 1% BSA) was used as secondary antibody.

### sSIRL-1 ELISA

To measure sSIRL-1 in cell supernatants, 96-wells flat-bottom MAXIsorp plates (Nunc) were coated overnight at 4°C with capture mAb 1A5 (5 μg/mL in PBS, 50 μL/well). After washing with PBS 0.05% (v/v) Tween-20, plates were blocked with 100 μL/well blocking buffer (1% (w/v) BSA, 3% (w/v) dry milk in PBS). Next, plates were washed and incubated with undiluted cell supernatants and the standard curve consisting of serially diluted sSIRL-1^ecto^ in PBS 1% BSA (50 μL/well), overnight at 4°C. The following day, plates were washed and incubated with biotinylated mAb 3D3 for 1 hour at RT, followed by washing and incubation with streptavidin poly-HRP (0.1 μg/mL; Sanquin) for 1 hour at RT. Finally, plates were washed and incubated with 100 μL/well TMB-substrate (Biolegend/ITK). The substrate reacted approximately 8 minutes, after which color development was stopped by adding 100 μL/well 1M H_2_SO_4_. Absorbance was measured at 450 nm on the CLARIOstar^®^ (BMG Labtech). Absorbance at 570 nm was used for background correction. All incubations, except coating of the capture antibody, were done on a shaker.

The same procedure was used to measure sSIRL-1 in plasma, serum, urine or sputum samples, with two exceptions: a different blocking buffer was used (3% (w/v) BSA in PBS), and samples were pre-incubated 15 minutes on a shaker with 20 μg/mL HAMA blocking reagent (Fitzgerald) before adding the samples to the ELISA plate, to prevent a-specific interactions.

We expressed the concentration of sSIRL-1 in pM rather than pg/mL, because endogenous sSIRL-1 may have a different molecular weight than the recombinant sSIRL-1^ecto^ that was used for the standard curve.

### Neutrophil stimulation with protease inhibitors

Neutrophils were isolated from peripheral blood using density gradient centrifugation on Ficoll (GE Healthcare). The neutrophil pellet was incubated with ammonium chloride buffer to lyse the erythrocytes. The remaining neutrophils were washed and suspended in RPMI 1640 containing 10% (v/v) FCS and 1% (v/v) PS. Neutrophils were then stimulated with 50-100 ng/mL tumor necrosis factor (TNF; Miltenyi) or 100 μg/mL of the Dectin-1 ligand Curdlan (Wako biochemical) in flatbottom plates (Nunc) at 37°C, with or without addition of protease inhibitors. After 2 to 4 hours, neutrophils were centrifuged 5 minutes at 500g to collect the supernatants. Cells were used for flow cytometry analysis. Supernatants were centrifuged once more for 30 minutes at 25000g at 4°C to remove cellular debris, followed by ELISA measurements.

### PLB-985 cell treatment with neutrophil serine proteases

The transduction and culture method of PLB-985 cells with SIRL-1 overexpression have been previously described (16). Cells were treated 2h at 37°C with neutrophil elastase (Elastin Products Company), cathepsin G (Biocentrum), or proteinase 3 (Elastin Products Company) (all 1 μM), followed by flow cytometry analysis.

### Flow cytometry

To determine SIRL-1 expression, cells were washed once with FACS buffer and stained with mAb 1A5 or 3F5 conjugated to AF647 or appropriate isotype controls (BD Biosciences) for 20 minutes at 4°C. Subsequently, the cells were washed twice with FACS buffer and finally taken up in FACS buffer with addition of 7AAD (BD Bioscience) for viability staining. Cells were analyzed using the BD FACS Canto II and FlowJo software (Treestar, Ashland, OR). For neutrophil activation experiments, neutrophils were gated as in Supplementary Fig 3. Neutrophil purity was always >90%, based on forward scatter (FSC) and sideward scatter (SSC).

### Statistical analysis

Statistical analyses were performed using GrapPad Prism software (version 8.3.0). For each graph, the statistical test and number of biological replicates (n) are described in the figure legends.

## RESULTS

### Development of sSIRL-1 ELISA

To develop a sSIRL-1 ELISA, we first produced recombinant His-tagged sSIRL-1 proteins: SIRL-1 ectodomain (sSIRL-1^ecto^) and VSTM1-v2, representing products of ectodomain shedding and alternative splicing, respectively (See Fig 1A for a schematic representation of the proteins, and Supplementary Fig 1A for the amino acid sequences). Both proteins were N-glycosylated, as shown by a shift in molecular weight after PNGase-F treatment and SDS-PAGE and Western blot analysis (Fig 1B, C). Deglycosylated VSTM1-v2, with a predicted MW of 21.6 kDa, had an apparent size of 37 kDa, which is consistent with the MW found in a previous study (22). Deglycosylated sSIRL-1^ecto^, with a predicted MW of 13.8 kDa, had an apparent size of 16 kDa. The size of the proteins was not affected by reduction with 2-mercaptoethanol, indicating the proteins did not form dimers with disulfide bonds.

For the ELISA, sSIRL-1^ecto^ was used as a standard, SIRL-1 mAb clone 1A5 as capture antibody and SIRL-1 mAb clone 3D3 as detection antibody. In a competition assay, clone 1A5 and 3D3 did not interfere with each other for binding to SIRL-1, and thus recognize different epitopes (Supplementary Fig 1B). The ELISA detected sSIRL-1^ecto^ and VSTM1-v2 equally well, with a lower limit of detection of 8 pM (Fig 1D). The ectodomain of the inhibitory receptor LAIR-1 (sLAIR-1^ecto^), which has 31% sequence identity with sSIRL-1^ecto^, was not detected in the ELISA, indicating specificity of the ELISA (Fig 1D). We spiked sSIRL-1^ecto^ into heparin plasmas that were sSIRL-1 negative in our ELISA. The spike was recovered in all plasma dilutions tested (Fig 1E), indicating that plasma is not interfering with the sSIRL-1 measurement. Next, we spiked sSIRL-1^ecto^ into human pooled serum (HPS) and subjected it to 10 freeze thaw cycles, or incubation at 37°C, 56°C or 65°C for 30 minutes or 1 hour. sSIRL-1 concentration was stable in all freeze thaw cycles but decreased after 30 minutes incubation at 65 °C (Supplementary Fig 1C). Taken together, we developed a sensitive and specific assay to measure sSIRL-1 protein concentration and showed that sSIRL-1 can be detected in presence of human plasma components and is stable during multiple freeze-thaw cycles.

### sSIRL-1 is present *in vivo* and increased in respiratory inflammation

To investigate the presence of sSIRL-1 protein *in vivo*, we used the ELISA to measure sSIRL-1 in plasma of healthy individuals, stratified per genotype of the rs612529T/C SNP. rs612529C associates with decreased SIRL-1 expression on monocytes, but not on granulocytes (14, 15). sSIRL-1 was detectable in only a small percentage of plasma samples from healthy individuals (8 out of 53, 15.1%) (Fig 2A). There was a tendency toward a lower percentage of samples with detectable sSIRL-1 in individuals with a rs612529 C allele, but these differences were not statistically significant (Fig 2B).

**Fig 2.**
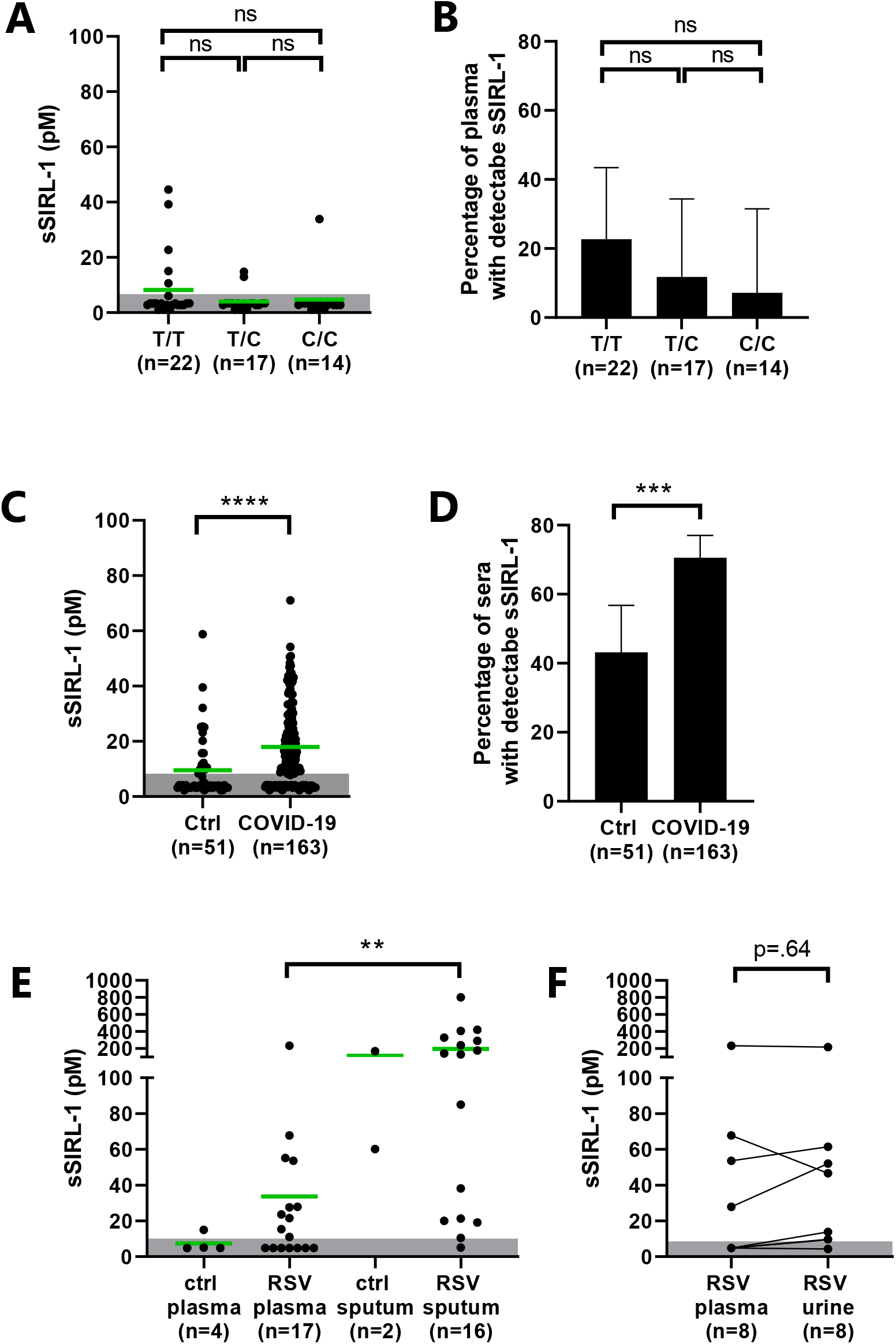
sSIRL-1 is increased in COVID-19 and RSV bronchiolitis patients. sSIRL-1 was measured by ELISA in plasma, urine and sputum. **(A-B)** sSIRL-1 concentration in plasma from 53 healthy donors, stratified per genotype of the rs612529 SNP (T/T, n=22; T/C, n=17; C/C, n=14). **(C-D)** sSIRL-1 concentration in serum of hospitalized COVID-19 patients (n=163) and control serum (n=51). **(E)** sSIRL-1 concentration in heparin plasma or sputum of mechanically ventilated infants with severe RSV bronchiolitis or mechanically ventilated infants without infection (controls). Control plasma, n=4; RSV plasma, n=17; control sputum, n=2; RSV sputum, n= 16. **(F)** sSIRL-1 concentration in paired plasma and urine samples of infants with severe RSV bronchiolitis (n=8) (**A, C, E, F)** Each dot represents one donor, the green bars represent the mean. The shaded area indicates the lower limit of detection (LLOD). Samples with undetectable sSIRL-1 were given a value of 0.5 x LLOD. Statistical differences were calculated using a Kruskal-Wallis test with Dunn’s correction (A, E), a Mann Whitney test (C), or a Wilcoxon test (F). **(B, D)** The percentage of samples with detectable sSIRL-1. The error bars indicate the 95% confidence interval, calculated with Wilson-Brown. The statistical differences were calculated using a Fisher’s exact test (B, D), in combination with Bonferroni correction for multiple testing (B). ** p ≤ .01, *** p ≤ .001. **** p ≤ .0001, ns = not significant.

To investigate the presence of sSIRL-1 in an inflammatory context, we measured sSIRL-1 in patients with COVID-19 or RSV bronchiolitis, which are both characterized by excessive neutrophil recruitment and activation (reviewed in (27, 28). We detected sSIRL-1 in approximately 70% of sera drawn from hospitalized adult COVID-19 patients, and the mean sSIRL-1 concentration was significantly higher than in control serum (Fig 2C-D). sSIRL-1 concentration in serum of COVID-19 patients was not affected by sex, age, nor the time since the start of symptoms or hospitalization (Supplementary Fig 2A, B, D, E). Within patients, sSIRL-1 concentration fluctuated in serum collected at two time points (Supplementary Fig 2C). Serum samples in this cohort were collected at non-standardized time points, and we had restricted availability to clinical data, thereby limiting further analysis on the correlation between sSIRL-1 concentration and disease activity.

We previously showed that RSV bronchiolitis patients have decreased SIRL-1 expression on sputum neutrophils compared to peripheral blood neutrophils (17, 29), which is in line with our hypothesis that activated neutrophils shed the ectodomain of SIRL-1. Indeed, in patients with severe RSV bronchiolitis, we detected sSIRL-1 in 15 out of 16 sputa, and the mean sSIRL-1 concentration in sputum was significantly increased compared to plasma (Fig 2E). We also detected sSIRL-1 in the urine of the RSV patients, in similar concentrations as in plasma (Fig 2F). In summary, we show that sSIRL-1 concentration is mostly undetectable in healthy individuals, but increased in hospitalized COVID-19 or severe RSV bronchiolitis patients.

### Activated neutrophils shed sSIRL-1 via proteolytic cleavage

To test if activated neutrophils indeed shed SIRL-1, we stimulated neutrophils from healthy controls *in vitro* with TNF or curdlan, with or without addition of a broad-spectrum protease inhibitor cocktail. We analyzed SIRL-1 expression on neutrophils by flow cytometry, and sSIRL-1 concentration in the supernatant by ELISA. In agreement with our previous work (16), the percentage of SIRL-1^+^ neutrophils decreased after activation (Fig 3A, B). Concomitantly, we detected sSIRL-1 in the supernatant (Fig 3C). Treatment with the protease inhibitor cocktail prevented this (Fig 3A-C), indicating that activated neutrophils release sSIRL-1 via proteolytic cleavage.

**Fig 3.**
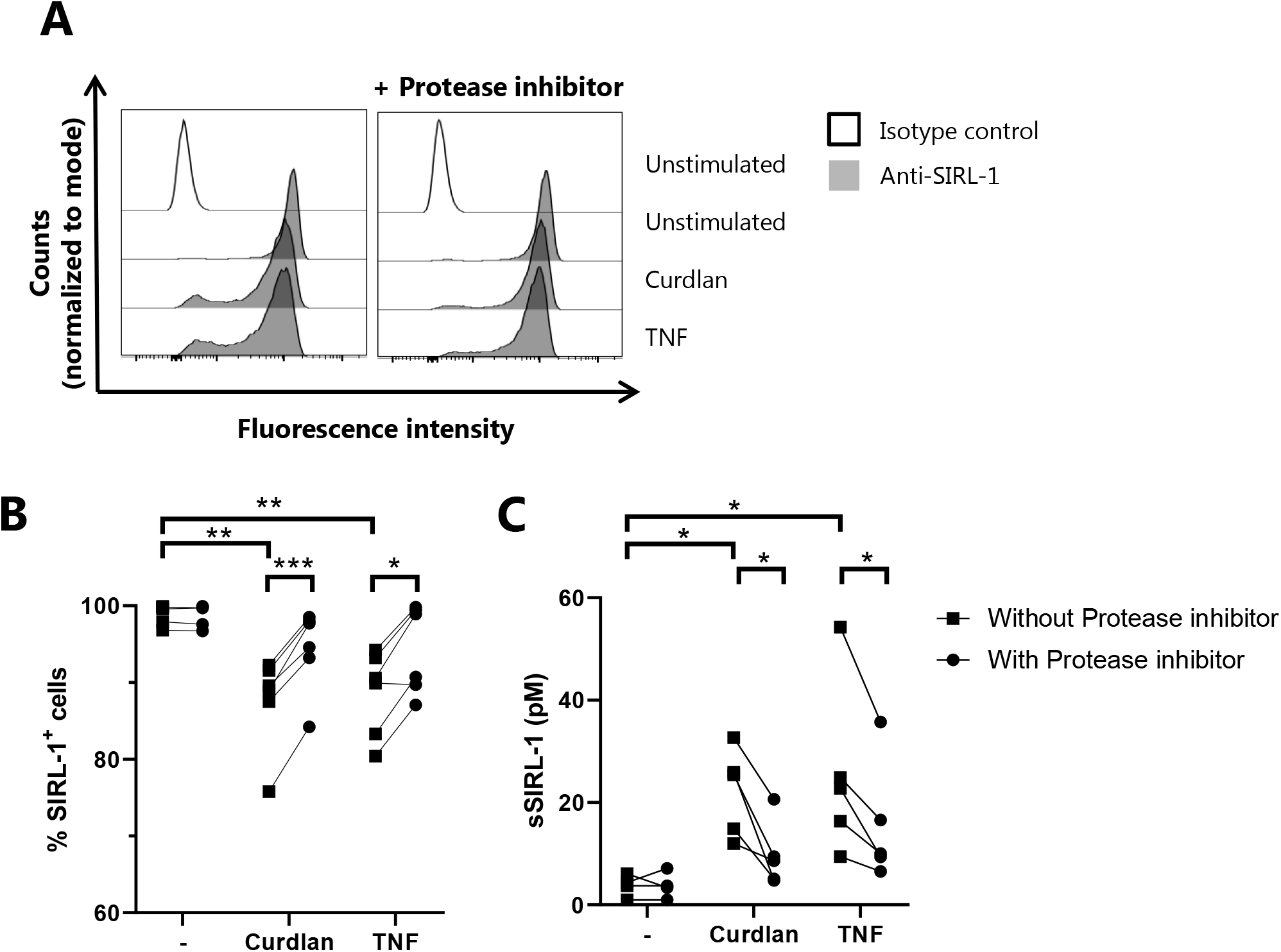
Activated neutrophils shed SIRL-1. Neutrophils were isolated from healthy donors and stimulated 4h at 37 °C with curdlan (100 μg/mL) or TNF (100 ng/mL), with or without addition of a protease inhibitor cocktail. **(A-B)** SIRL-1 expression on neutrophils was analyzed by flow cytometry. Shown are representative histograms of the fluorescence intensity of cells stained with a SIRL-1 antibody (clone 1A5) (closed histogram) or an isotype control (open histogram) (A) and the quantification of the percentage of SIRL-1^+^ cells (B). Each symbol represents one donor, n=6. **(C)** Supernatants from stimulated neutrophils were analyzed by sSIRL-1 ELISA. Each symbol represents one donor, n=5. Statistical significance was determined using a two-way ANOVA with Geisser-Greenhouse correction and Holm-Sidak’s multiple comparison test (B-C), * p ≤ .05, ** p ≤ .01, *** p ≤ .001.

### sSIRL-1 is shed by proteinase 3

To examine which protease cleaves SIRL-1, we activated neutrophils with TNF in combination with inhibitors against major classes of proteases: pepstatin A for aspartic proteases, E64 for cysteine proteases, GM6001 for metalloproteases, leupeptin for serine and cysteine proteases, and aprotinin for serine proteases. Treatment with 10 μM aprotinin resulted in a small but significant increase in the percentage of SIRL-1-expressing cells after TNF stimulation, suggesting that SIRL-1 is shed by a serine protease (Fig 4A). Thus, we further investigated SIRL-1 cleavage by the serine proteases that are predominantly secreted from activated neutrophils; cathepsin G, neutrophil elastase, and proteinase 3 (PR3) (30). Treatment of SIRL-1-overexpressing PLB-985 cells with PR3 resulted in a modest but consistent decrease in SIRL-1 expression, whereas cathepsin G and elastase did not affect SIRL-1 expression on these cells (Fig 4B-C). To confirm that PR3 cleaves SIRL-1, we activated neutrophils with TNF in presence of a specific PR3 inhibitor. Indeed, the PR3 inhibitor partially prevented the loss of SIRL-1 expression on TNF-activated neutrophils in all donors tested (Fig 4D-E). Finally, treatment of sSIRL-1^ecto^ and VSTM1-v2 with PR3 in a purified system resulted in proteolytic cleavage as assessed by SDS-PAGE and Western Blot (Fig 4F). Thus, we show that PR3 cleaves SIRL-1.

**Fig 4.**
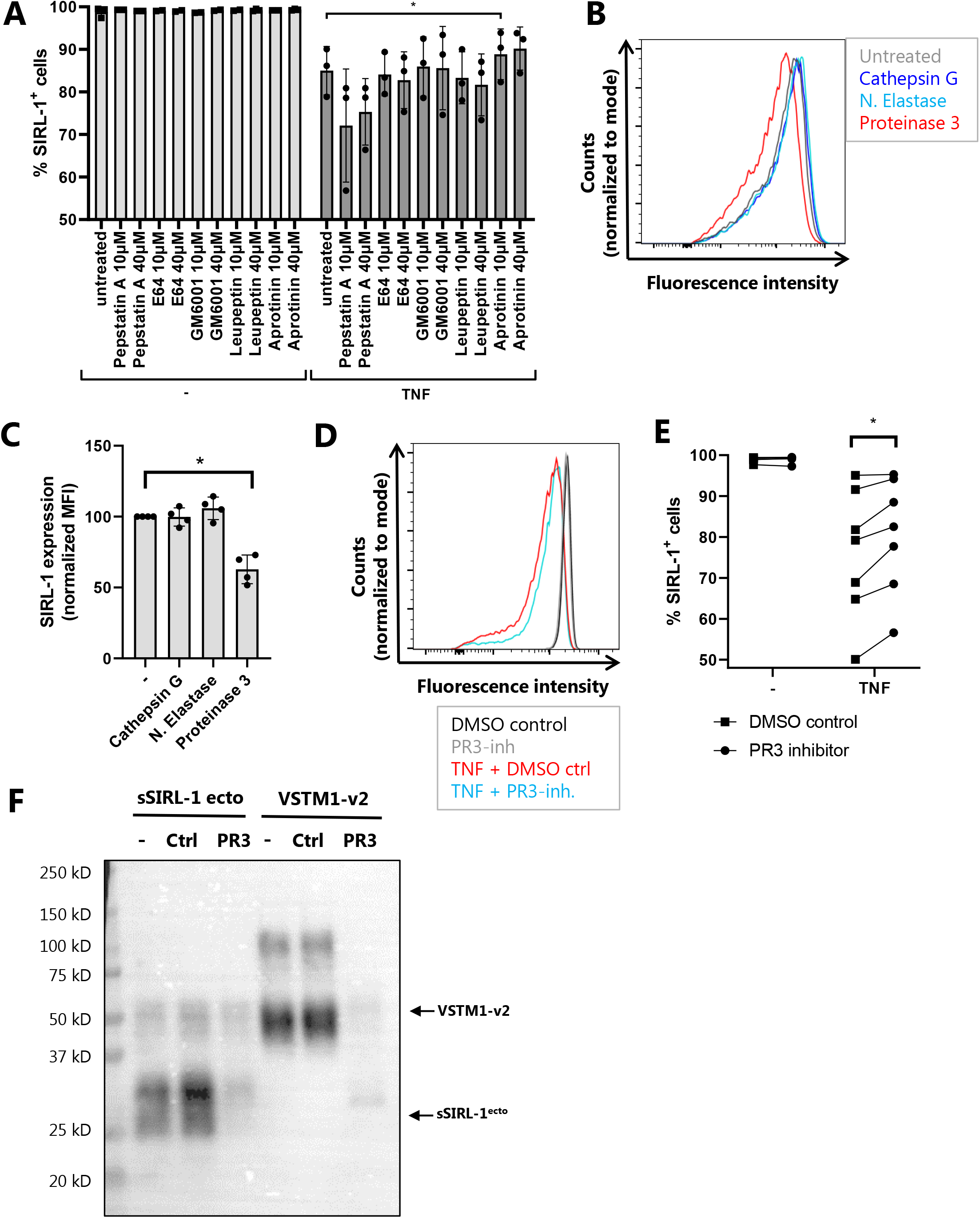
SIRL-1 is cleaved by PR3. **(A, D, E)** Neutrophils were isolated from healthy donors and stimulated 2h at 37 °C with TNF (50 ng/mL), with or without addition of inhibitors against major protease classes (A; 10-40 μM) or a PR3 inhibitor (D-E; 10 μM). Cells were stained with a SIRL-1 antibody (clone 3F5) and analyzed by flow cytometry. **(A)** The bars indicate the percentage of SIRL-1^+^ cells (mean ± SD), each symbol represents a donor, n=3. **(D)** Representative histograms of the fluorescence intensity of cells stained with a SIRL-1 antibody and **(E)** the quantification of the percentage of SIRL-1^+^ cells (each symbol with connected line represents a donor), n=7. **(B-C)** PLB-985 cells with SIRL-1 overexpression were treated 2h with neutrophil elastase, cathepsin G, or proteinase 3 (all 1 μM), followed by flow cytometry analysis (n=4). Shown are representative histograms of the fluorescence intensity of cells stained with a SIRL-1 antibody (clone 3F5) (B) and the quantification of the median fluorescent intensity (MFI), normalized to the MFI of untreated cells (C; mean ± SD, each symbol represents one experiment). **(F)** sSIRL-1^ecto^ and VSTM1-v2 were left untreated (-) or treated with buffer control (ctrl) or PR3 for 3h at 25 °C, and analyzed by SDS-PAGE and Western Blot. The membrane was stained with a SIRL-1 antibody (clone 1A5). The arrows indicate the position of VSTM1-v2 or sSIRL-1^ecto^. One representative experiment of n=3 is shown. Statistical significance was determined using a mixed-effects model with Dunnett’s multiple comparisons test (A; TNF treatment alone was compared to TNF treatment with each of the protease inhibitors), a 1-way ANOVA with Dunnett’s multiple comparison test (C) or a mixed-effects model with Sidak’s multiple comparisons test (E), all with Geisser-Greenhouse correction, * p ≤ .05.

### *S. aureus* protein Eap inhibits sSIRL-1 shedding by neutrophils

SIRL-1 is activated by α-type phenol-soluble modulins (PSMs) from *Staphylococci* (19). *S. aureus* is known for its large repertoire of secreted factors that can influence the host immune response, including inhibitors against neutrophil proteases. Among these, *S. aureus* produces extracellular adherence protein (Eap) and the homologues EapH1 and EapH2, which specifically inhibit neutrophil serine proteases, including PR3 (24). We therefore questioned whether Eap can inhibit shedding of SIRL-1 by neutrophils. Remarkably, treatment with Eap almost completely prevented the loss of membrane-expressed SIRL-1 after TNF stimulation of neutrophils (Fig 5A, B). A similar trend was seen after treatment with EapH2, whereas treatment with EapH1 had no effect on membrane SIRL-1 expression. In PLB-985 cells, Eap and EapH1 both inhibited SIRL-1 cleavage by exogenous PR3 (Fig 5C). Together, these data indicate that Eap can inhibit proteolytic cleavage of SIRL-1.

**Fig 5.**
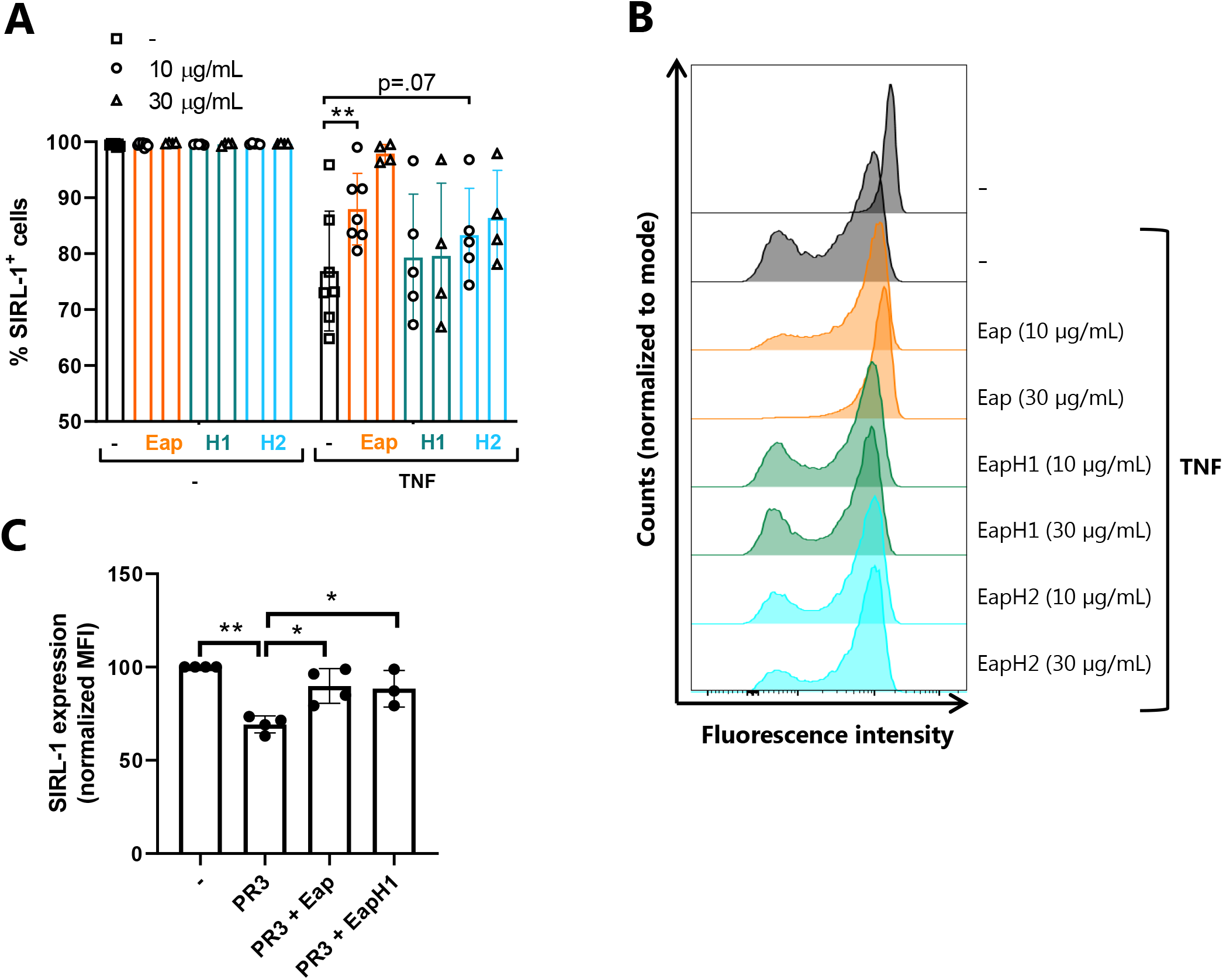
*S. aureus* protein Eap inhibits SIRL-1 shedding. **(A-B)** Neutrophils were isolated from healthy donors and stimulated for 2h at 37 °C with TNF (50 ng/mL), with or without addition of 10-30 μg/mL Eap, EapH1, or EapH2 (indicated with H1 or H2, respectively). Cells were stained with a SIRL-1 mAb (clone 3F5) and analyzed by flow cytometry, n=4-6. (A) The percentage of SIRL-1^+^ cells (mean ± SD), each symbol represents a donor. (B) Representative histograms of the fluorescence intensity of cells stained with a SIRL-1 antibody. (C) PLB-985 cells with SIRL-1 overexpression were treated 2h with 1 μM proteinase 3, with or without addition of Eap or EapH1 (30 μg/mL). SIRL-1 expression was analyzed by flow cytometry. The bars indicate the percentage of SIRL-1^+^ cells (mean ± SD), each symbol represents an experiment, n=3-4. Statistical significance was determined using a mixed-effects model with Dunnett’s multiple comparisons test (A) or Holm-Sidak’s multiple comparisons test (C), both with Geisser-Greenhouse correction, * p ≤ .05, ** p ≤ .01.

## DISCUSSION

Here, we developed an ELISA to measure sSIRL-1 concentration (Fig 1) and are the first to show that sSIRL-1 protein is present *in vivo* (Fig 2).

sSIRL-1 concentration was increased in sputum of RSV bronchiolitis patients compared to plasma, suggesting release of sSIRL-1 at the site of infection (Fig 2E). sSIRL-1 was also detectable in sputum of control infants, which can be explained by previous observations that sputum neutrophils of these control patients are activated (17), possibly due to mechanical ventilation (31). From the site of infection, sSIRL-1 may leak into the circulation, as there was a trend of increased sSIRL-1 in RSV patient plasma compared to control plasma, although this comparison was limited by the low number of control plasmas (Fig 2E). Similarly, sSIRL-1 concentration was increased in serum of hospitalized COVID-19 patients compared to control serum (Fig 2C). Lastly, sSIRL-1 was also detectable in urine of RSV patients (Fig 2F), indicating that, similar to soluble LAIR-1 (26), sSIRL-1 is cleared by the kidneys.

Several studies have indicated a role of neutrophils in the pathophysiology of COVID-19 (reviewed in (28). Using a machine learning algorithm on a broad panel of inflammatory markers, neutrophil-related markers predicted critical illness of COVID-19 patients most strongly (32). These markers included resistin, lipocalin-2 and hepatocyte growth factor, which are all released by degranulating neutrophils. Similarly, PR3 was highly present in sputum of COVID-19 patients (33), and its concentration predicted COVID-19 disease severity (34). Due to restricted availability of data and the collection of serum at non-standardized time points, we could not correlate sSIRL-1 concentration to COVID-19 disease progression, clinical parameters or inflammatory markers in our cohort. Future studies will have to clarify if sSIRL-1 may serve as a biomarker to predict COVID-19 disease severity, or other diseases characterized by high neutrophil activation.

We found that SIRL-1 is cleaved from activated neutrophils by PR3 (Fig 3-4). In addition to secretion of soluble PR3, activated neutrophils can express PR3 on the plasma membrane on a subpopulation of cells (35). Membrane-bound PR3 (mPR3) has been suggested to bind to the plasma membrane via CD177, other membrane-expressed proteins, or directly to the lipid bilayer (36–38). We compared shedding of SIRL-1 on FACS-sorted CD177^+^ versus CD177^-^ neutrophils and found that CD177 expression did not affect sSIRL-1 shedding (unpublished observations). Still, we showed that SIRL-1 surface expression decreased in a subpopulation of neutrophils after activation *in vitro* (Fig 3A, 4D, 5B), which may reflect cells with high mPR3 expression. On the other hand, SIRL-1 expression was homogeneously low on neutrophils in sputum of RSV bronchiolitis patients (17, 29). Hence, the relative contribution of soluble PR3 and mPR3 to SIRL-1 shedding remains to be determined.

Finally, we show that shedding of SIRL-1 by activated neutrophils was prevented by Eap, a neutrophil serine protease inhibitor secreted by *S. aureus* (Fig 5). The homologues EapH1 and EapH2 were less effective in preventing SIRL-1 shedding from activated neutrophils. In a previous study, using short peptide substrates, EapH1 and EapH2 also had a lower capacity to inhibit PR3 than Eap (Eap, K_i_ = 0.23 nM; EapH1, K_i_ = 1.0 nM; EapH2, K_i_ = 21 nM) (24) (Fig 5A, B). Alternatively, EapH1 and EapH2 may differ in their ability to inhibit mPR3, as mPR3 has been shown to be more resistant to PR3 inhibitors than soluble PR (39). The latter finding may also explain the partial effectiveness of the PR3 inhibitor used in this study, and the differential abilities of leupeptin and aprotinin to inhibit SIRL-1 shedding (Fig 4A, 5A).

In addition to ectodomain shedding, sSIRL-1 may derive from the splice variant VSTM1-v2 (22, 40). VSTM1-v2 mRNA expression was increased in PBMCs of Rheumatoid Arthritis patients compared to controls (40). However, it remains to be determined whether endogenous VSTM1-v2 protein is present *in vivo*. Both forms of sSIRL-1 were recognized by our ELISA (Fig 1D), thus not allowing for discrimination between these forms. Further research into VSTM1-v2 protein would benefit from the development of an antibody that recognizes the intracellular tail of SIRL-1, which is present in VSTM1-v2 but not in shed sSIRL-1.

Currently, we can only speculate on the function of SIRL-1 shedding. We previously argued that IIRs that are constitutively expressed on a cell form a threshold to prevent unnecessary immune activation. Some of these threshold receptors, so-called disinhibition receptors, are downregulated after an activating stimulus surpasses the initial threshold, to facilitate subsequent cellular activation (2). SIRL-1 is such an disinhibition receptor, based on its constitutive high expression on monocytes and granulocytes in peripheral blood and lung (12, 14, 15) and downregulation during inflammation (17, 29). We thus propose that the function of SIRL-1 shedding is to rapidly remove SIRL-1 to facilitate for a strong anti-microbial response, once the threshold for activation is surpassed.

Of course, the inhibitory function of SIRL-1 and hence the effect of SIRL-1 shedding also depends on expression of its ligands. We found that SIRL-1 is activated by S100 proteins (20), cathelicidin LL-37, and PSMs from Staphylococci (19). In inflammatory conditions with local tissue damage and DAMP release, neutrophil released LL-37 and S100 proteins may act mostly on newly incoming neutrophils with high SIRL-1 expression, to signal to these cells that no further immune activation is required.

PSMs are produced by both pathogenic and harmless Staphylococci (41). SIRL-1 ligation by PSMs may be beneficial for the host, for example by facilitating tolerance of resting neutrophils to harmless Staphylococci such as *S. epidermidis*, but still allowing for full neutrophil activation once SIRL-1 is shed in an inflammatory context. Interestingly, *S. aureus*, the most pathogenic member of the Staphylococcus family, is unique in its secretion of Eap proteins (24). *S. aureus* may use Eap to inhibit SIRL-1 shedding to benefit from the preserved inhibitory function of membrane-expressed SIRL-1. Of particular interest is a recent study showing that *S. aureus* also requires Eap to prevent degradation of PSMα3 by neutrophil serine proteases (42), indicating that Eap potentially preserves expression of SIRL-1 as well as its ligands.

In conclusion, we measured increased sSIRL-1 concentration in COVID-19 and RSV bronchiolitis patients and provided mechanistic insight into the loss of SIRL-1 from activated neutrophils. Future studies will have to further elucidate the functional implications of sSIRL-1 release, and its potential use as a biomarker.

## Acknowledgements

We thank Brice Korkmaz and Adam Lesner for generously providing the PR3 inhibitor, Anouk van Haperen for asistance with cloning and production of recombinant sSIRL-1 proteins, Jop Wattel for assistance with the development of the sSIRL-1 ELISA, Prof. Wang De Yun for subject recruitment for the SSIC samples, Tamara Brouwers for assistance with neutrophil stimulation assays, Maaike Ressing for the suggestion to test the effect of *S. aureus* protease inhibitors on SIRL-1 shedding, and Femke van Wijk for valuable suggestions on the manuscript. This work was supported by a Vici grant from the Netherlands Organization for Scientific Research (NWO, grant no. 91815608) and a ZonMW grant (grant no. 10430012010024).

## Authorship

LM and HR conceptualized the study. HR, GW, and DG performed the experiments. HR and GW analyzed the data. ED, SN, HL and KR were responsible for the design and sample collection of the COVID-19 patient cohort, SB and LB for the RSV patient cohort, and AA and OR for the rs612529 cohort. PM was involved in the development and validation of SIRL-1 mAb clone 3F5. SR and DS provided Eap, EapH1, and EapH2, and contributed to the design and interpretation of the experiments with these proteins. LM, LB and SR supervised the study. HR wrote the initial draft of the manuscript. All authors contributed to the writing and editing of the final manuscript. LM and SN acquired funding for the study.

## Declaration of interest

None

## Notes

### Competing Interest Statement

The authors have declared no competing interest.

