## Supplementary files for "Soluble signal inhibitory receptor on leukocytes-1 is released from activated neutrophils by proteinase 3 cleavage"

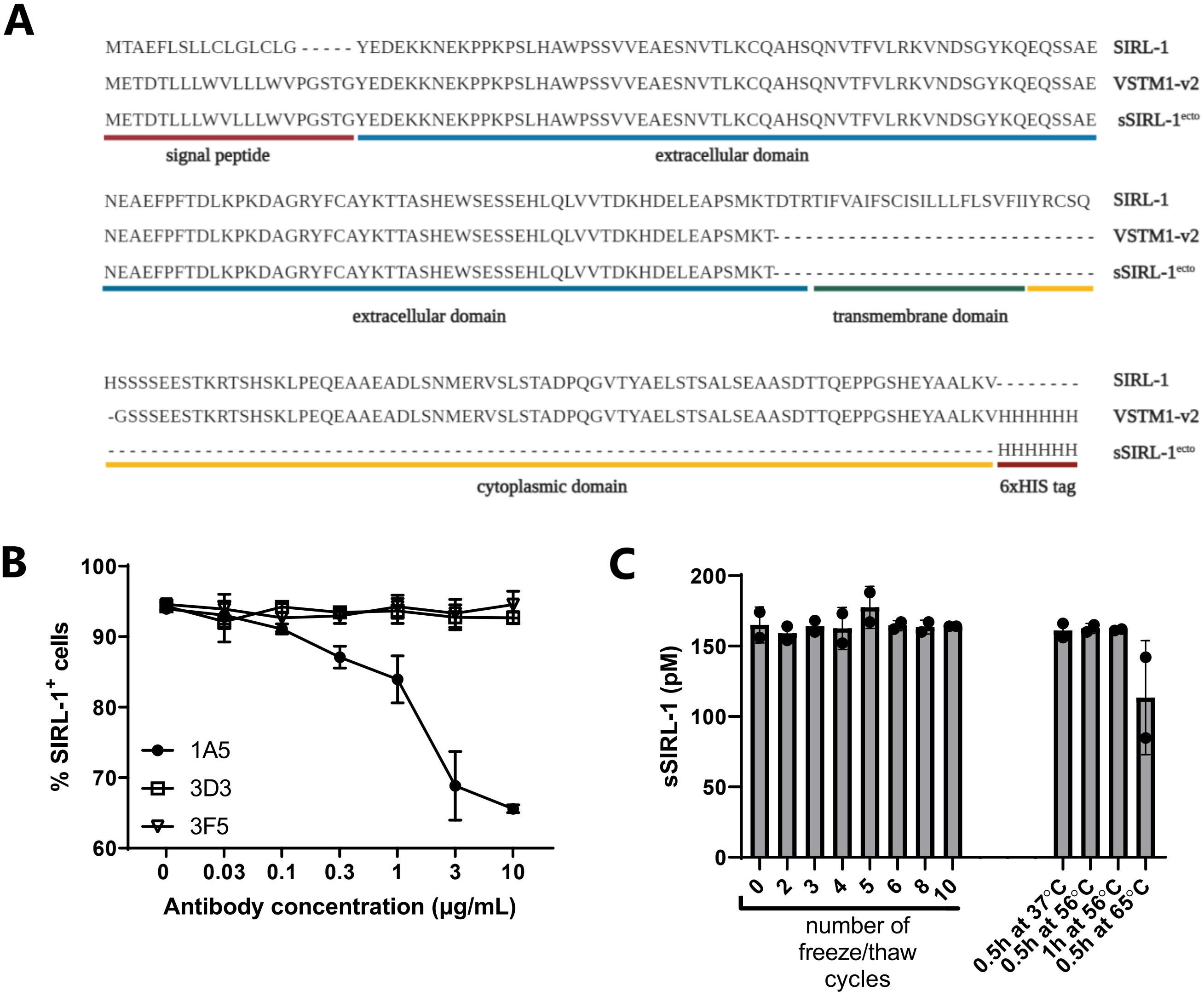


**Supplementary Fig 1. Development of sSIRL-1 ELISA**
**(A)** Amino acid sequence of membrane-expressed SIRL-1, and recombinant VSTM1-v2 and sSIRL-1$^{ecto}$. Horizontal colored lines indicate the signal peptide, extracellular domain, transmembrane domain, cytoplasmic domain and His-tag. For sSIRL-1$^{ecto}$, the sequence of the ectodomain of SIRL-1 minus three amino acids was used. **(B)** PBMCs were pre-incubated for 2 hours at 37°C with 0.03 – 10 µg/mL of indicated anti-SIRL-1 mAb clones, followed by staining with anti-SIRL-1 clone 1A5-AF647 and flow cytometry analysis. The graph shows the percentage of SIRL-1 expressing cells, n=1, mean ± SD from technical duplicates. **(C)** Human pooled serum was spiked with sSIRL-1$^{ecto}$ and subjected to a maximum of ten freeze-thaw cycles, 0.5 or 1h incubation at 56 °C, or 0.5h incubation at 65 °C. sSIRL-1 concentration was measured by ELISA, n=2.

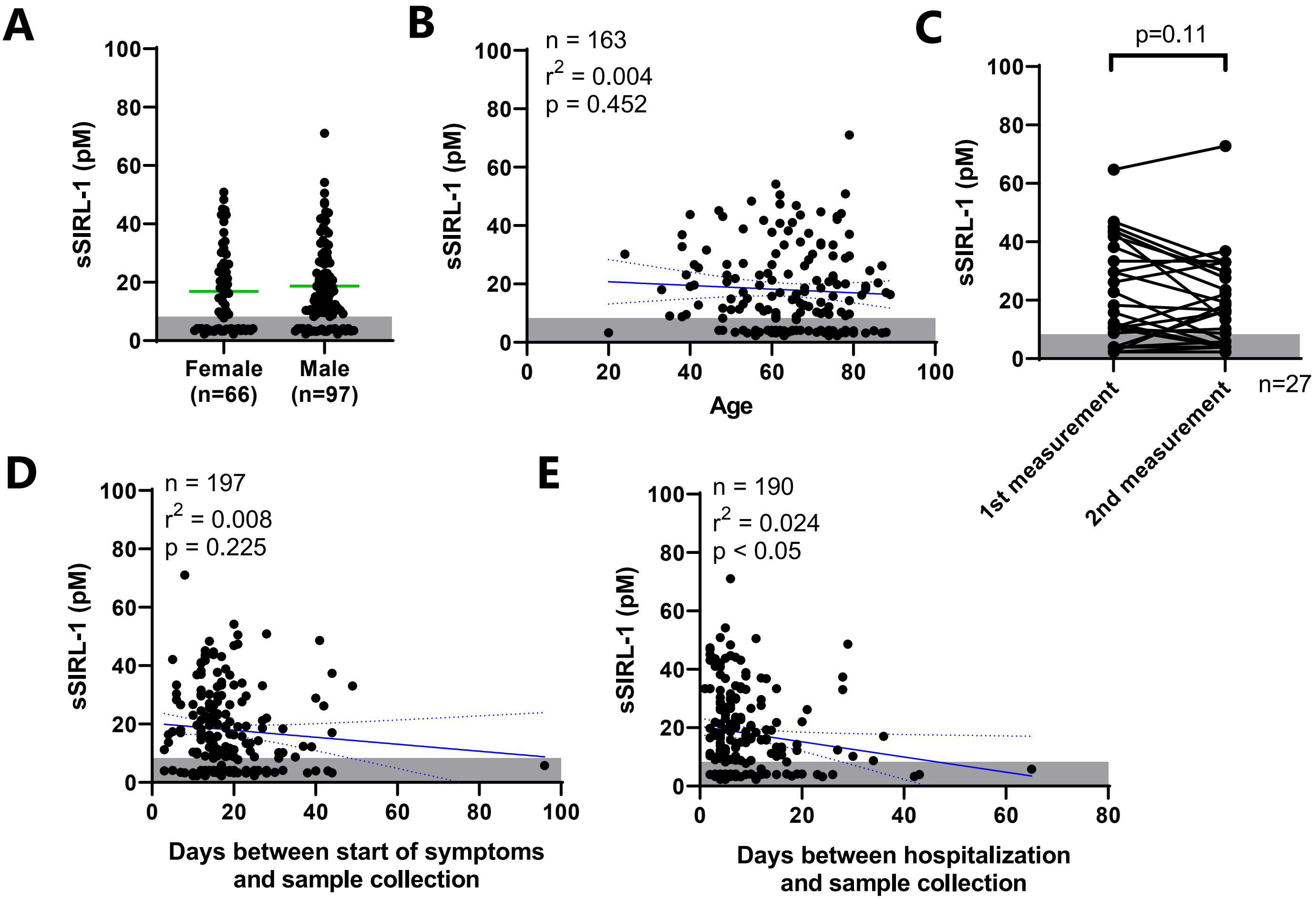


**Supplementary Fig 2. sSIRL-1 concentration in hospitalized COVID-19 patients**

sSIRL-1 was measured by ELISA in serum of hospitalized COVID-19 patients at one (n=138) or two (n=27) time points at various intervals, and correlated to sex (A), age (B), number of days between start of symptoms and sample collection (D) or number of days between hospitalization and sample collection (E). Graphs show only the first measurement time point per donor (A-B), compare the first and second measurement time point within the same patient (C), or show all measurement time points (D-E). Each dot represents one donor, the green bars represent the means. The shaded area indicates the lower limit of detection (LLOD), and samples with undetectable sSIRL-1 were given a value of 0.5 x LLOD. Correlations were calculated using simple linear regression, the blue lines represent the best fit line with 95% confidence interval. In C, statistical significance was determined using a Wilcoxon matched-pairs test.

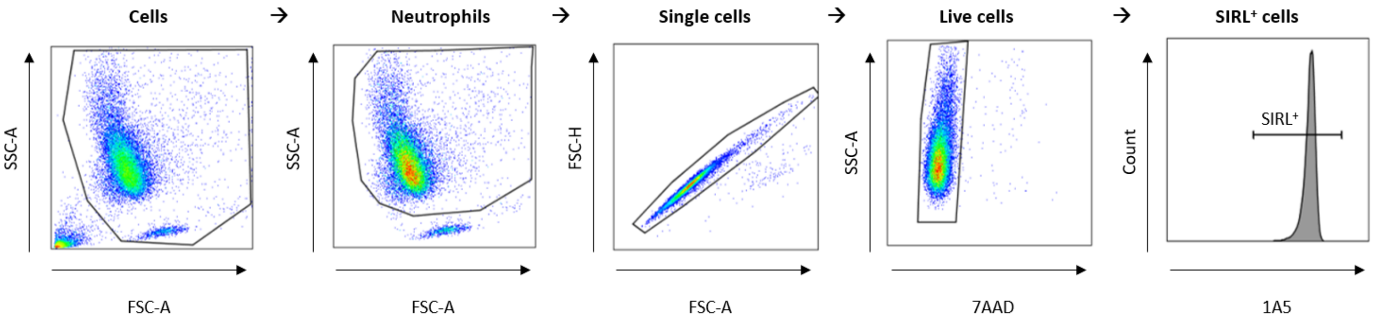


**Supplementary Fig 3. Gating strategy of *in vitro* stimulated neutrophils**

Neutrophils were isolated by ficoll gradient centrifugation, stimulated *in vitro* and analyzed by flow cytometry. Single neutrophils were gated using forward scatter (FSC) and sideward scatter (SSC). Viable cells were selected by negative 7AAD staining, followed by selection of SIRL-1$^+$ cells.

| Recombinant protein | cDNA sequence |
| --- | --- |
| sSIRL-1$^{ecto}$ | ATGGAGACAGACACACTCCTGCTATGGGTACTGCTGCTCTGGGTTCCAGGTTCCACTGGCTACGAAGATGAGAAAAAGAATGAGAAACCG<br>CCCAAGCCCTCCCTCCACGCCTGGCCCAGCTCGGTGGTTGAAGCCGAGAGCAATGTGACCCTGAAGTGTCAGGCTCATTCCCAGAATGTGA<br>CATTTGTGCTGCGCAAGGTGAACGACTCTGGGTACAAGCAGGAACAGAGCTCGGCAGAAAACGAAGCTGAATTCCCCTTCACGGACCTGA<br>AGCCTAAGGATGCTGGGAGGTACTTTTGTGCCTACAAGACAACAGCCTCCCATGAGTGGTCAGAAAGCAGTGAACACTTGCAGCTGGTGG<br>TCACAGATAAACACGATGAACTTGAAGCTCCCTCAATGAAAACACACCACCATCATCACCACTAATAA |
| VSTM1-v2 | ATGGAGACAGACACACTCCTGCTATGGGTACTGCTGCTCTGGGTTCCAGGTTCCACTGGCTACGAAGATGAGAAAAAGAATGAGAAACCG<br>CCCAAGCCCTCCCTCCACGCCTGGCCCAGCTCGGTGGTTGAAGCCGAGAGCAATGTGACCCTGAAGTGTCAGGCTCATTCCCAGAATGTGA<br>CATTTGTGCTGCGCAAGGTGAACGACTCTGGGTACAAGCAGGAACAGAGCTCGGCAGAAAACGAAGCTGAATTCCCCTTCACGGACCTGA<br>AGCCTAAGGATGCTGGGAGGTACTTTTGTGCCTACAAGACAACAGCCTCCCATGAGTGGTCAGAAAGCAGTGAACACTTGCAGCTGGTGG<br>TCACAGATAAACACGATGAACTTGAAGCTCCCTCAATGAAAACAGGTTCATCATCTGAGGAATCCACCAAGAGAACCAGCCATTCCAAACTT<br>CCGGAGCAGGAGGCTGCCGAGGCAGATTTATCCAATATGGAAAGGGTATCTCTCTCGACGGCAGACCCCCAAGGAGTGACCTATGCTGAG<br>CTAAGCACCAGCGCCCTGTCTGAGGCAGCTTCAGACACCACCCAGGAGCCCCCAGGATCTCATGAATATGCGGCACTGAAAGTGCACCACC<br>ATCATCACCACTAATAA |

**Supplementary Table 1. cDNA sequence of recombinant sSIRL-1 proteins**

The table indicates the cDNA sequence for recombinant sSIRL-1$^{ecto}$ and VSTM1-v2.
